# Ribosome profiling reveals the translational landscape and allele-specific translation efficiency in rice

**DOI:** 10.1101/2022.02.22.481533

**Authors:** Xi-Tong Zhu, Run Zhou, Jian Che, Yu-Yu Zheng, Muhammad Tahir ul Qamar, Jia-Wu Feng, Jianwei Zhang, Junxiang Gao, Ling-Ling Chen

**Affiliations:** National Key Laboratory of Crop Genetic Improvement, College of Informatics, Huazhong Agricultural University, Wuhan 430070, China; State Key Laboratory for Conservation and Utilisation of Subtropical Agro-bioresources, College of Life Science and Technology, Guangxi University, Nanning 530004, China

**Keywords:** Ribo-seq, translational landscape, uORFs, lncORFs, allele-specific translation efficiency, rice (*Oryza sativa*)

## Abstract

Translational regulation is a critical step in the process of gene expression and governs the synthesis of proteins from mRNAs. Many studies have revealed the translational regulation in plants in response to various environmental stimuli. However, there has been no comprehensive landscape of translational regulation and allele-specific translation efficiency in multiple tissues of plants, especially in rice, a main staple crop feeding nearly half of the world population. Here, we used RNA-seq and Ribo-seq data to analyze the transcriptome and translatome of an elite hybrid rice SY63 and its parental varieties ZS97 and MH63. The results revealed that gene expression patterns varied more significantly between tissues than between varieties at both transcriptional and translational levels. Besides, we identified 3,392 upstream open reading frames (uORFs), and most of the uORF-containing genes were enriched for transcription factors. Only 668 long non-coding RNAs could be translated into peptides. Finally, we discovered numerous genes with allele-specific translation efficiency in SY63, and further demonstrated that some *cis*-regulatory elements (secondary structures of mRNAs and the binding of miRNAs) may contribute to allelic divergence in translation efficiency. Overall, our findings may improve the understanding of translational regulation in rice and provide information for the molecular basis of breading research.

## Introduction

Understanding the molecular basis for phenotypic variations is essential in eukaryotic biology research, especially the variations in gene expression. Transcription and translation are two critical steps in the process of gene expression (Crick, 1970). Over the past few decades, transcriptomic studies have described the gene expression profiles of a wide variety of species in animals (Cardoso-Moreira *et al*., 2019) and plants (Klepikova and Penin, 2019), and revealed the complexity of gene regulatory networks and evolution of gene expression patterns in different species. However, proteomics studies are relatively lagging behind largely due to limitation by the relatively low throughput of mass spectrometry (Noor *et al*., 2021). Ribosome profiling (Ribo-seq), a technique for deep-sequencing of ribosome-protected RNA fragments (RPFs) (Ingolia *et al*., 2012), enables both global monitoring of translation process *in vivo* and quantification of translated open reading frames (ORFs) in RNA, and thus may be used as a proxy of protein synthesis (Brar and Weissman, 2015). Moreover, when combined with corresponding mRNA sequencing (RNA-seq), Ribo-seq can be used to investigate the dynamics of translation efficiency on a genome-wide scale (Ingolia *et al*., 2009). The assessment of translation efficiency can indirectly decipher the effect of extensive translational regulation that results in poor correlations between RNA abundance and protein level (Wang *et al*., 2020).

Unlike mammals, plants generally exhibit broad variations within their genomes and correspondingly complex translational profiles due to several ancient whole genome duplication events and the following chromosomal rearrangements (Clark and Donoghue, 2018). Understanding the role of translational changes on gene expression thus is more challenging but important and many researchers attempted to describe it in plants, particularly in *Arabidopsis* (Juntawong *et al*., 2014) and maize (Lei *et al*., 2015). More recently, a study in rice revealed the effect of nitrogen application after abrupt drought-flood alternation on translation and revealed that a fraction of genes were up- or down-regulated in either transcriptome or translatome (Xiong *et al*., 2020). Although these studies have provided valuable insights into the translational regulation of some important crops, there is still a lack of comprehensive understanding of the translation status in multiple tissues of plants. In addition, although research on translational regulation in plants is still at the initial stage, researchers have begun to investigate the divergence in the translation efficiency of alleles in hybrid yeast (Artieri and Fraser, 2014; Muzzey *et al*., 2014) and mice (Hou *et al*., 2015). For example, Hou et al. first detected over one thousand genes with allele-specific translation efficiency (ASTE) in hybrid mice and demonstrated the effects of *cis*-regulatory elements (Hou *et al*., 2015). They found that many common *cis*-regulatory elements such as the binding of microRNAs (miRNAs) are not associated with the ASTE of genes. However, it remains undetermined whether this finding is also true in plants.

In this study, to gain more insights into ASTE and provide a comprehensive translational regulation profile in rice (*Oryza sativa*), we conducted RNA-seq and Ribo-seq experiments for the leaves, panicles and roots of three Asian rice varieties (*Oryza sativa* ssp. *xian/indica*), including Minghui 63 (MH63), Zhenshan 97 (ZS97), and their elite hybrid Shanyou 63 (SY63). We compared the gene expression between the transcriptional and translational levels and investigated the differences in translational efficiency (TE) among different tissues. As a result, thousands of actively translated upstream open reading frames (uORFs) and hundreds of active ORFs were detected from mRNAs and long non-coding RNAs (lncRNAs), respectively, which enables a full characterization of translational regulation in rice. Besides, based on the identified genes with ASTE, we demonstrated that some *cis-* regulatory elements on mRNAs influence the TE in alleles, which helps to extend the related knowledge in animals to plants.

## Results

### Data quality and characteristics of ribosome profiling in rice

To obtain a comprehensive translational landscape of multiple rice varieties and tissues, we carried out Ribo-seq for three rice varieties: the male parent MH63, female parent ZS97, and their elite hybrid SY63. Three tissues, including leaves and roots at seedling stage and panicles, were collected for each variety with two biological replicates per tissue and variety (Figure 1A). Both RNA-seq and Ribo-seq were carried out on the same materials, and the RNA-seq data have been recently published in our previous work (Zhou *et al*., 2021). Here, Ribo-seq and RNA-seq data were mapped to the gap-free genome MH63RS3 (Song *et al*., 2021) to determine the data quality and the overall landscape of translational regulation. The mean Pearson correlation coefficient between two biological replicates was 0.92, indicating a good reproducibility of our Ribo-seq data (Figure S1A).

**Figure 1.**
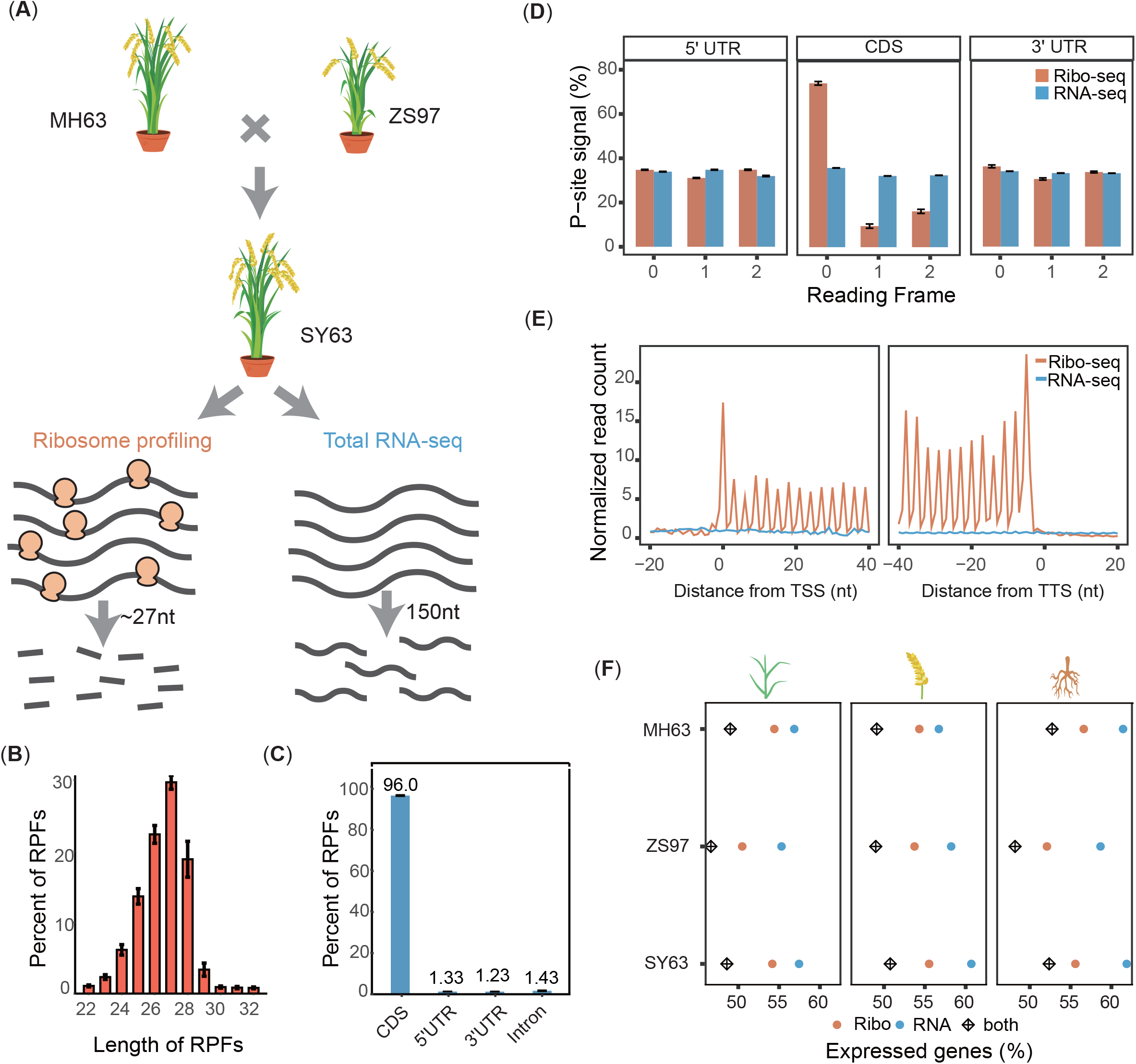
Deep sequencing-based global quantification of parents and hybrids. (A) Brief overview of the experiment design. Detailed protocols for the Ribo-seq and RNA-seq experiments can be found in the Methods. (B) Length distribution of Ribo-seq ribosome-protected fragment (RPFs), with peak at 27 nt. (C) The proportion of RPFs within annotated genes, ∼96% of which are located in the CDS region. (D) Reading frame of optimally mapped Ribo-seq reads and RNA-seq reads within annotated genes. (E) Distribution of optimally mapped reads along the CDS within each codon. Each read was represented by a specific P-site position depending on its fragment length. TSS, translation start site; TTS, translation termination site. (F) The proportion of expressed genes (TPM>0.1) at translatome (orange), transcriptome (blue) and both levels across tissues and varieties. Data in b-e were calculated for each replicate and aggregated together. RPFs with size from 25-31nt were used in d and e. Error bars display ± *s. e. m*.

Ribo-seq data possess several unique features that can be utilized to monitor the dynamic translation process and determine the quality of ribosome profiling libraries (Ingolia *et al*., 2009; Liu *et al*., 2013; Juntawong *et al*., 2014; Lei *et al*., 2015; Wu *et al*., 2019). The dominant length of RPFs reported in animals and maize (Lei *et al*., 2015) is 28–30 nt. However, in rice, we observed that the vast majority of RPFs are 26–28nt in length, particularly RPFs of 27nt (Figure 1B). This unique feature was also observed in a previous study, in which a high-quality rice ribosome footprint library was constructed (Yang *et al*., 2020). In total, 96.0% of RPFs could be mapped to annotated coding region (CDS), while the remaining RPFs were almost equally distributed in 5’UTR, 3’URT and intron (Figure 1C). In ribosome, peptidyl elongation occurs in the P-site (Lauria *et al*., 2018), and the estimated P-site offset varied depending on the read length. For example, the reads of 27nt corresponded to the P-site offset of 11nt (Figure S1C). Most importantly, reading frame analysis of the mapped RPFs revealed that more than 70% of RPFs were accumulated in the first frame of CDS (Figure 1D), while no enrichment of RPFs was found in 5’UTR and 3’UTR. Moreover, corresponding to codon triplets, a clear three-nucleotide (3-nt) periodicity could be detected when using the first nucleotide of the P-site as the position of ribosome footprint on mRNAs, while no 3-nt periodicity or frame preference was found in RNA-seq data (Figure 1E). There was a higher density of reads near the end region of mRNAs, which could be attributed to the usage of rare codons (Figure S1D) (Wang and Roossinck, 2006). Finally, among the 39,046 non-transposable element genes in MH63RS3 genome, 50%–60% of them could be transcribed and translated (Figure 1F), which is consistent with a previous report (Zhao *et al*., 2017). Besides, once transcribed, an average of 85% genes would recruit the ribosome complex for translation (Table S1), indicating the protein coding function of these genes. These results suggested that our Ribo-seq data for all three varieties and tissues were of high quality (Dunn *et al*., 2013; Fields *et al*., 2015) and valid for further analysis.

### Expression changes at transcriptional and translational levels

Based on the transcriptome and translatome data, we first investigated the global differences in gene expression patterns among different varieties and tissues. There were high correlations (∼0.95; Figure S2A) between the normalized expression values (TPM) generated from 150 bp paired-end (PE) reads and simulative 30 bp single-end (SE) reads, indicating that the length of RNA-seq reads would cause little bias in the analysis results. This enabled a proper comparison between the two expression levels and the computation of the TE. Next, we computed the variance of expression at both levels in order to measure the effects of transcriptional or translational regulation on gene expression (Wang *et al*., 2020). Various types of post-transcriptional modifications can regulate gene expression by modulating translation efficiency, leading to higher expression variance in translatome (Csárdi *et al*., 2015; Weinberg *et al*., 2016). Consistently, we found that the expression variances in translatome were higher than those in transcriptome for all varieties and tissues (Figure 2A, 2B, S2B). On average, the Ribo-seq variances for the leaf, panicle and root were ∼0.733, ∼0.858 and ∼0.793, which were about 8.39%–25.3% higher than those in RNA-seq (Figure 2B). Genome-wide analysis of TE variation range showed that the panicle had the broadest TE range (∼167 fold), but no consistent pattern was observed for the three varieties (Figure S2C). In contrast, the TE distribution displayed a clear tissue-specific pattern: the leaf had higher TEs than the other two tissues (Figure 2C), indicating that tissue-specific regulations at both transcriptional and translational levels together contribute to the difference in TE among tissues. To check whether some specific genes are strongly influenced due to the translational regulation (Lei *et al*., 2015; Wang *et al*., 2020) in rice, we conducted a GO enrichment analysis and found functional divergence between highly and poorly translated genes. For example, in MH63 leaf, the top 5% efficiently translated genes were mainly enriched in the interaction with environment, such as “response to stimulus (GO:0050896, *P*=2.12e-12)” and “response to stress (GO:0006950, *P*=1.76e-11)” (Figure 2D). However, low TE genes were particularly enriched in very basic cell life process like “DNA metabolic process (GO:0006259, *P*=1.13e-06)” (Figure 2D, Data S2). Besides, although there were relatively high correlations of gene expression between the two levels (Figure S3), high TE genes did not necessarily have high transcriptional abundance (Figure S2D, S2E). These results suggested that TE can be used as an indicator to study the gene expression regulation in rice.

**Figure 2.**
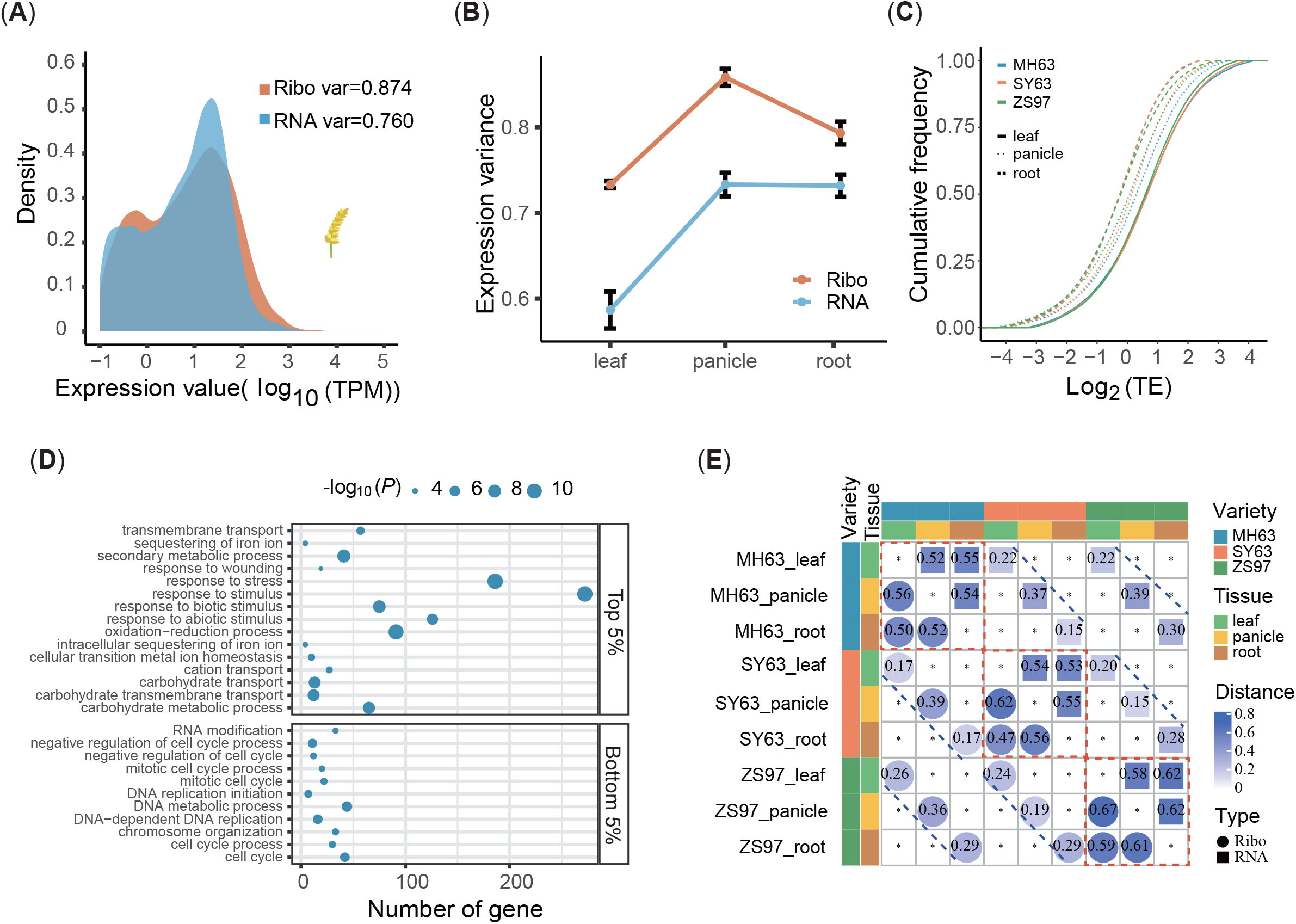
Gene expression divergence between transcriptome and translatome. (A) The distribution of expression value at translational level in panicle of SY63, compared with that at transcriptional level. The expression variance at translational level is greater than that of transcriptome. (B) The expression variances were calculated for both translatome and transcriptome levels across tissues and varieties. Error bars indicate ± *s. e. m*. (C) Cumulative distribution of TE for protein-coding genes expressed across tissues and varieties. (D) GO enrichment results for top 5% and bottom 5% TE genes in MH63 leaf. (E) Euclidean distance of expression profile at the translational and transcriptional levels across tissues and varieties.

Moreover, we calculated the divergence of expression profiles between or within the transcriptional and translational level for all varieties and tissues. As shown in Figure 2E, the square and circle represent the gene expression distance at the two levels. The data in the three red dashed boxes and blue dashed line represent the expression distances of RNA-seq and Ribo-seq in different tissues of the same variety and the same tissue of different varieties, respectively. It could be seen that the expression distances in the red dashed box were significantly larger than those on the blue dashed line at both transcriptional level and translational level. In general, the inter-tissue expression distances in the same variety were much larger than the inter-variety distances in the same tissue, suggesting that the same tissue shares similar transcriptional and translational regulation mechanisms among different varieties. It could be observed that the transcription distance or translation distance of the same tissue was roughly equal between varieties. These results implied that gene expression is regulated by both transcription and translation rather than dominantly by either of them.

### Regulation of uORFs on translation in rice

Although a previous study has identified thousands of uORFs in rice, this study was mainly focused on the effect of abrupt drought-flood alternation (Xiong *et al*., 2020). Here, we comprehensively characterized the uORFs among three varieties and different tissues. A total of 2,902, 2,474 and 2,912 uORFs were identified in MH63, ZS97 and SY63, respectively (Table S2, Data S3). Although each variety harbored hundreds of unique active uORFs, about two thirds of uORFs were shared by all the three varieties (Figure 3A), indicating that these uORFs are necessary and might participate in important cell processes. As expected, the length of active uORFs was shorter than that of main ORFs (mORFs), with a median value of 96nt (Figure 3B, S4A). The core function of uORFs is to suppress the translation of mORFs in animals (Calvo *et al*., 2009; Ingolia, 2014) and plants such as maize (Lei *et al*., 2015). However, it has been reported that genes with one uORF have similar TEs to the genes without uORFs, and when the number of uORFs increases to two or above, the uORFs could improve the TEs of their mORFs in rice young leaves (Xiong *et al*., 2020). Here, we found a tissue-specific pattern of the regulatory effect of uORFs on mORFs. In the root, with increasing number of uORFs, the TE of mORFs would be slightly decreased (Figure 3C). However, for the leaf and panicle, the number of uORFs had almost no effect on mORFs (Figure S4B). We also calculated the variation coefficients of TEs to evaluate the effects of uORFs. Interestingly, the variation coefficients of TEs for genes without uORFs were significantly higher than those with one or more uORFs for all varieties and tissues (Figure 3C, S4C). This was consistent with another function of uORFs recently found in *Drosophila melanogaster* (Patraquim *et al*., 2020), which reported that uORFs can buffer the translation of mORFs, leading to a lower dispersion of TEs.

**Figure 3.**
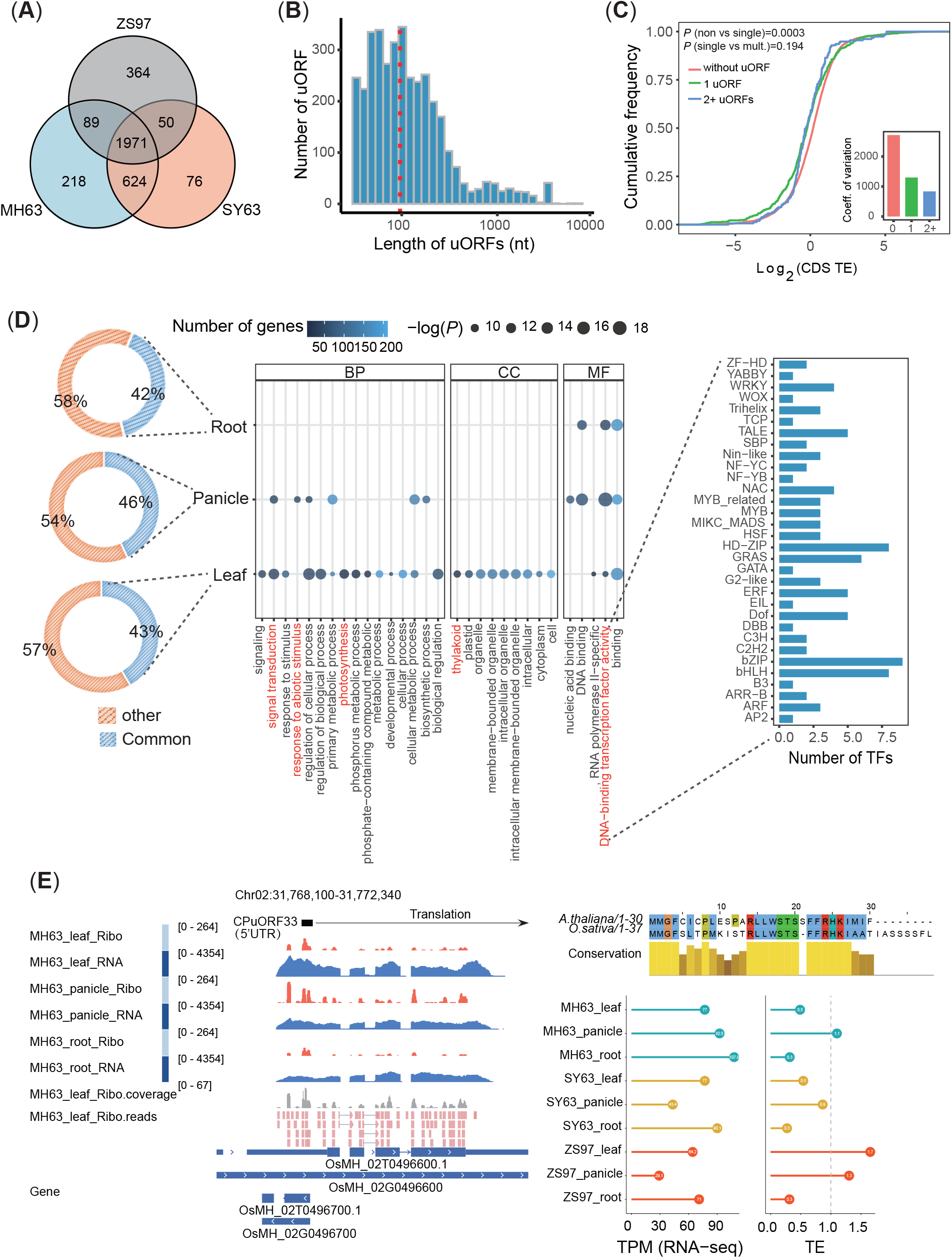
uORF-mediated translational regulation. (A) Three-way Venn diagram showing the numbers of uORFs shared in three varieties. (B) Length distribution of the translated uORFs, with the median size of 92nt (read dash line). (C) Cumulative distribution of CDS TEs in main ORFs and coefficient of variation grouped by their number of uORFs in the root of SY63. Kolmogorov-Smirnov test p-values are labeled. (D) The GO terms and transcription factor categories of the uORF-contained genes shared by the three varieties in different tissues. The left panel shows the proportion of uORF shared by the three varieties across tissues. The right panel shows the GO terms of the uORF-contained genes shared by the three varieties across tissues. (E) The gene *OsMH_02G0496600*, homology of *ATHB1* in *Arabidopsis*, contains the same uORF called CPuORF33 which also inhibits the translation of the main ORF. The Ribo-seq and RNA-seq signals from leaf, panicle and root of MH63 are shown on the left panel (only tissues from MH63 were shown). The raw reads from MH63 leaf are also listed to explain that these signals are from CPuORF33, not the anti-sense gene *OsMH_02G0496700*. The protein sequence comparison of CPuORF33 in *Arabidopsis* and rice are shown on the top right panel and the expression level as well as TEs of the main ORF in *OsMH_02G0496600* are shown on the bottom right panel.

Considering that a large proportion of uORFs were shared by all samples (Figure 3D), we conducted a GO enrichment analysis to understand the potential function of these uORF-containing genes. As shown in Figure 3D, while some GO terms were tissue-specific, “DNA-binding transcription factor activity (GO:0003700, *P*_max_=1.94e-04)” was significantly enriched for all three tissues, indicating that these genes play certain roles as transcription factors (TFs). By searching plant TF databases, we found that many uORF-containing genes harbor various TF DNA-binding domains as shown in the right panel of Figure 3D. Some of these genes may be vital to plant development. For example, *OsMH_02G0496600* is the homologue of *ATHB1* (*AT3G01470*) in *Arabidopsis* that encodes a homeodomain-leucine zipper (HD-Zip) domain. Overexpression of *ATHBI* caused deleterious phenotypes such as serrated leaves in *Arabidopsis*, and an uORF (CPuORF33) can repress the translation of this TF via a ribosome stalling mechanism in aerial tissues of *Arabidopsis* (Ribone *et al*., 2017). A similar DNA sequence of this uORF was identified in rice, and these two uORFs had a relatively high identity to each other (66.7%, Figure 3E). The expression profile showed that the mORF of this gene was expressed in all tissues (Figure 3E), yet the TE was below 1 in most tissues, indicating the suppression of this mORF by the presence of CPuORF33 homologue. Besides, the homologous gene (*OsMH_09G0287500*) of the heat shock factor *AtTBF1* (*AT4G36990*) (Pajerowska-Mukhtar *et al*., 2012) also harbors an uORF, indicating that the recognition of uORFs in this study is accurate (Data S3). Taken together, these identified uORFs are indispensable regulatory elements for gene expression and may be a valuable resource for further experimental studies in rice.

### Translation ability of lncRNAs in rice

To understand the translation ability of widely distributed lncRNAs reported in our previous work (Zhou *et al*., 2021), we re-identified the lncRNAs in MH63RS3 genome with the same pipeline and predicted their coding potential. In total, only 4.95% (668) of all lncRNAs with one or more active ORFs could be translated (Figure 4A, Data S4), suggesting that the identification of a majority of lncRNAs in our previous study is reliable and the translation of lncRNAs is not universal in rice. Although these active ORFs had a longer median length (237nt) than uORFs, they were still significantly shorter than ORFs from protein-coding genes (Figure 4B, S4A). Five types of lncRNAs were previously classified and we found that 60.3% of active ORFs were derived from long intergenic noncoding RNAs (lincRNAs), followed by long noncoding natural antisense transcripts (lncNATs; 23.4%; Figure 4C). Considering the poor conservation of lncRNAs among varieties (Zhou *et al*., 2021), and the relatively large number of shared active translational ORFs among varieties and tissues (Figure S5A), we speculate that these lncRNAs may have similar functions to protein-coding genes.

**Figure 4.**
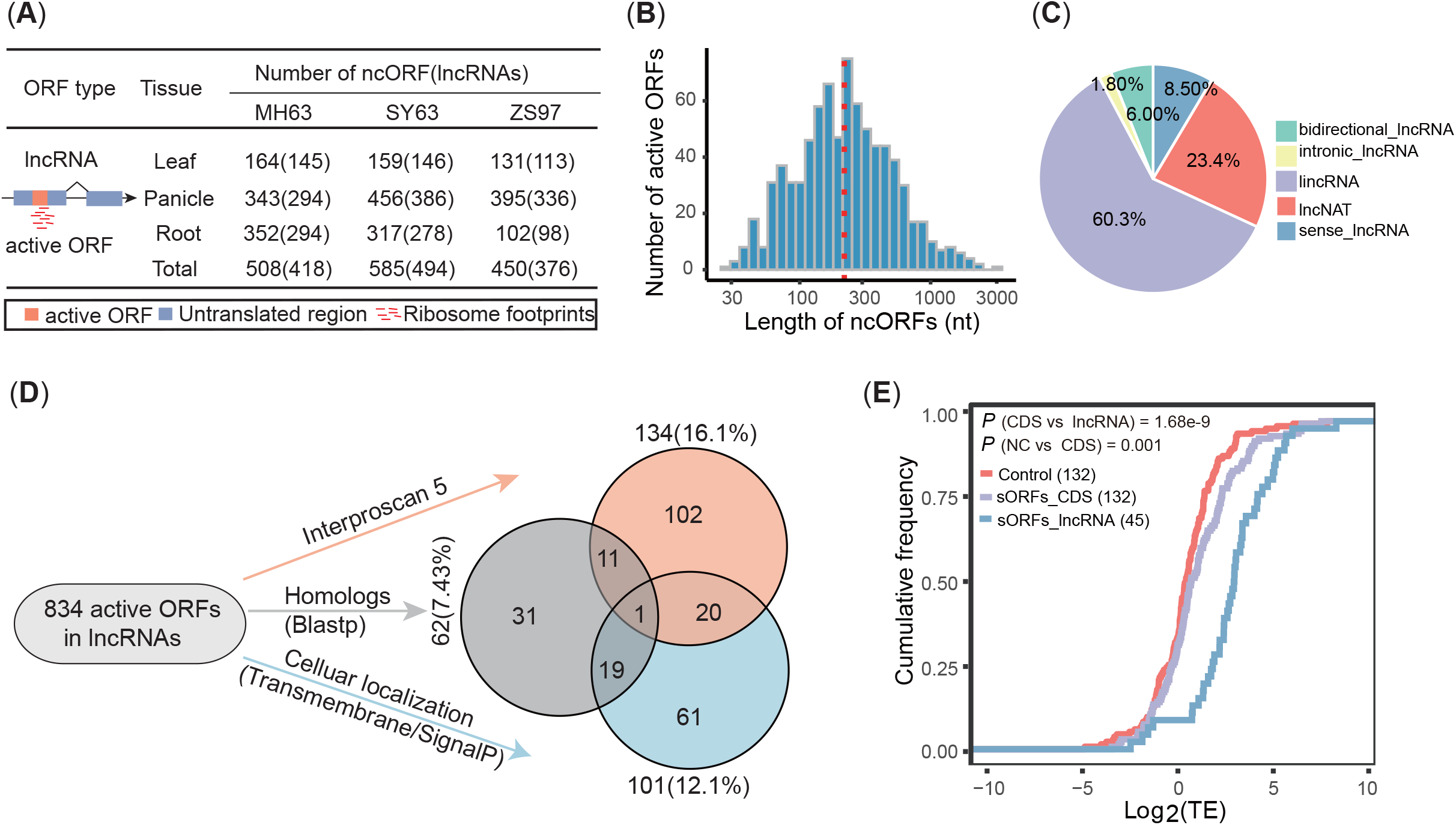
Translation of lncRNA genes. (A) The number of actively translated ORFs in lncRNAs identified in each tissue and variety. (B) Length distribution of active ORFs, with a median size of 237nt (read dash line). (C) Pie chart of lncRNA categories containing active ORFs. (D) Functional annotation of active ORFs. Venn chart showing the overlapped number of active ORFs identified with three different strategies. See Methods. (E) Cumulative distribution of TEs in lncRNA with sORFs, compared with protein-coding genes with sORFs in leaf of MH63. The TEs of randomly selected protein-coding genes whose CDS length were greater than 300nt in corresponding sample was used as a control. Kolmogorov-Smirnov test p-values among different kind of genes were labeled.

To validate this speculation, we conducted *de novo* functional annotation for a total of 834 active ORFs in lncRNAs from three aspects (Figure 4D, Data S4). First, we searched multiple protein databases using InterProScan 5 (Jones *et al*., 2014) and found that among the 834 active ORFs, 16.1% (134) contained one or more conserved domains. Second, we found that 62 (7.43%) active ORFs had high sequence similarities to known proteins through searching against all the annotated small proteins (shorter than 100 aa) in MH63RS3 genome. Third, considering that the small proteins can function as signal peptides in a wide range of plant species (Matsubayashi, 2014), we predicted the transmembrane topology and signal peptide structure for these active ORFs, finding that 101 (12.1%) might be located on the membrane (82) or be secreted (31). The majority of active ORFs (589; 70.6%) had no known functions (Data S4), which poses a great challenge for dissecting their possible roles during rice development.

Moreover, considering the relatively short length of active ORFs (Figure S5B), we next attempted to determine whether there are certain differences between active ORFs and the mRNAs containing small ORFs (sORFs; CDS size shorter than 300nt), such as translational efficiency. Surprisingly, we found that 243, 292 and 206 lncRNAs, and 300, 286 and 259 mRNAs could be translated into small peptides in MH63, SY63 and ZS97, respectively (Table S3). Then, we compared the TEs of sORFs from mRNAs and lncRNAs. The results showed that active ORFs had higher TEs than mRNAs (Figure 4E, S5C). Interestingly, the translation of these sORFs in lncRNAs tended to recruit rare codons (Figure S6B), and we speculated that the higher TEs could be more attributed to the shorter length of these sORFs (Figure S6A) (Zhao *et al*., 2017).

### Identification of genes with allele-specific translation efficiency (ASTE) in hybrid rice

In addition to the above findings, our data can also be used to determine the TE divergence between allelic genes in hybrid SY63 to identify the *cis*-regulatory elements responsible for such divergence. Since both allelic genes in hybrid are influenced by the same *trans*-acting factors (such as TFs and miRNAs), the observed TE divergence in alleles may be caused by *cis-*regulatory elements in mRNAs, such as GC content, secondary structure and miRNA binding site (McManus *et al*., 2014). To test this possibility, it is crucial to assign sequenced reads to definite parent alleles. We designed a pipeline to obtain such reads and identified the genes with ASTE (Figure 5A, Methods and Materials). Phased reads were obtained using PP2PG (Feng *et al*., 2021). The average error rate for Ribo-seq and RNA-seq data was 2.10% and 1.13%, respectively, and the separation rate in Ribo-seq was as low as 1.49% on average due to their short lengths (∼30nt); however, for RNA-seq reads, about 14.90% of them could be phased (Table S4, Data S5). Then, we selected 14,659 genes containing SNPs within their CDS region from all 1:1 orthologs between MH63 and ZS97 to identify the genes with significant ASTE. Under the cut-off *FDR* ≤ 0.05 and |*log*_2_(*Fold change*)| ≥ 1, we identified 420 genes with ASTE, including 81, 36 and 358 in the leaf, panicle and root of SY63, respectively (Data S6, S7). Only a few ASTE genes were shared by all three tissues (Figure S7A). The right panel of Figure 5A shows a representative example: the *ONAC096* (*OsMH_07G0033800*/*LOC_Os07g04510*) gene, which belongs to the NAC transcription factor superfamily (Puranik *et al*., 2012), was reported to mediate abscisic acid (ABA)-induced leaf senescence and improve the grain yield (Kang *et al*., 2019). Consistent with these findings, the expression level of this gene was high in the root but low in the panicle in all three varieties (Figure S7B). However, although there was no significant difference between the two alleles at the transcriptional level in the root of SY63, the Ribo-seq signal from the MH63 allele was much stronger than that from ZS97, indicating a strong allele specificity at the translational level (Figure S7C).

**Figure 5.**
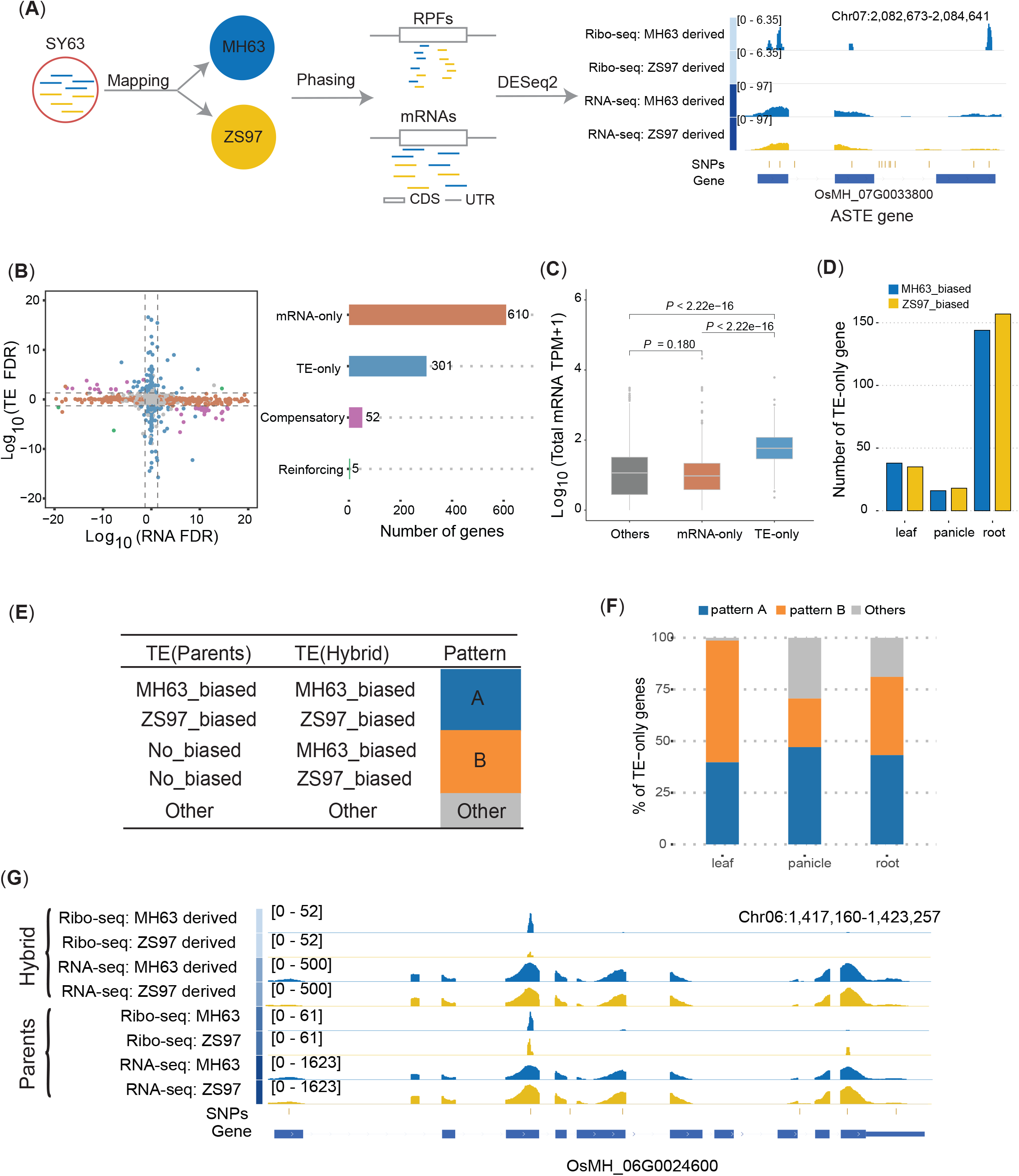
Characteristics of genes with ASTE. (A) The left panel is the pipeline used to identify genes with ASTE, which includes reads mapping, phasing with PP2PG and difference detection by DESeq2. See methods for details. The right panel is an example of gene with ASTE. Gene *OsMH_07G0033800* were shown with Ribo-seq and RNA-seq signals from both alleles. Four out of five SNPs within CDS region are covered by Ribo-seq reads in MH63 allele, indicating a reliable divergency in allelic TE. (B) Relationships between TE and RNA abundance in genes with allelic expression bias in root tissue. False discovery rate (FDR) for each gene represented by each point in left panel was adjusted according to its fold change (FC): −*log*_10_(*FDR*) *if FC*(*ZS*97/*MH*63) > 0; *log*_10_(*FDR*) *if FC*(*ZS*97/*MH*63) < 0 at mRNA or TE level. The right panel represented the number of different kind of genes in the left panel. (C) Comparison of total mRNA expression level among three kinds of genes in root tissues. Wilcoxson test p-values were labeled. Number of mRNA-only and TE-only genes were displayed in (B) and other gene was randomly selected from those are not in the four categories with equal number to mRNA-only genes. (D) The number of MH63 biased or ZS97 biased TE-only genes in three tissues of SY63. (E) Two major regulatory patterns exhibited in TE-only genes. Pattern A means the changes at TE levels are same in parents and hybrid, while the TE level is biased towards to one allele in pattern B. (F) Number of TE-only genes with different patterns in three tissues of SY63. (G) RNA-seq and Ribo-seq signals for gene *OsSPX-MFS3* (*OsMH_06G0024600* / *LOC_Os06g03860*) in hybrid (first four tracks) and parents (next four tracks), which shows switched pattern B and is reported to involve in maintaining phosphate homeostasis in rice.

Previous studies have reported multiple patterns of allelic regulation at both transcriptional and translational levels in yeast (Artieri and Fraser, 2014) and mice(Hou *et al*., 2015). Thus, we also explored this phenomenon in rice by taking the root as an example since it had the largest number of ASTE genes (358). Following the procedures in previous studies (Hou *et al*., 2015), all genes with allelic bias were classified into four categories (Data S8). Among these genes, 610 showed significant allelic specificity only in mRNA abundance (hereafter referred to as mRNA-only genes). On the contrary, 301 genes had significant allelic specificity only in TE (hereafter referred to as TE-only genes). The mRNA-only genes mean that the two alleles are only different at the transcriptional level, but their TEs are not significantly different. Similarly, the TE-only genes mean that the two alleles are only different at TE, but there is no significant difference at the transcriptional level. Other 57 genes had allelic specificity in both mRNA abundance and TE. As shown in Figure 5B, a positive coordinate value indicates that the gene was biased to ZS97, while a negative coordinate value suggests that the gene was biased to MH63. Among these 57 genes, 52 were in the second and fourth quadrant, exhibiting a compensatory effect of transcription and translation, which were called compensatory genes. These results suggested that in hybrid rice, compensatory regulation of transcription and translation is important, which may be utilized by hybrids to maintain the equal protein levels of the two alleles. Other five genes were in the first and third quadrants, showing a reinforcing effect of transcription and translation process, and thus were designated as reinforcing genes.

Furthermore, we compared the transcriptional level between mRNA-only and TE-only genes in all three tissues. The results showed that TE-only genes had significantly higher total mRNA abundance (Figure 5C, S7D, S7E), demonstrating the importance of translational regulation for highly expressed alleles. Moreover, TE-only genes were involved in various processes, while very few enriched functions were found for mRNA-only genes (Figure S9A, Data S9). Last, we categorized the TE-only genes into either MH63 biased or ZS97 biased types, and found that in all three tissues of SY63, these two types of genes were highly similar in number (Figure 5D).

To understand the potential role of translational regulation in heterosis, we further analyzed the TE-biased pattern of all TE-only genes among the parents and the hybrid variety with the similar pipeline above (Figure 5A, Materials and Methods). The TE of the two allele of a TE-only gene in the hybrid had three possibilities: MH63-biased, ZS97-biased, and no-biased (Figure 5E), and it was the similar case for a TE-only gene between the two parents. In total, the TE bias of the hybrid and parents had nine possible combinations, among which four accounted for a large proportion of the TE-only genes (Data S10), and thus were named as pattern A and pattern B (Figure 5E). In pattern A, the hybrid and parents had the same TE bias, while in pattern B, the TE turned from no-biased to biased towards one allele. Pattern A comprised 43.3% of the total TE-only genes in average (Figure 5F). Among these genes, 87.2% could be functionally annotated, which was significantly higher than the ratio of randomly selected genes (fisher exact test, *P* = 0.009482). Among those genes with similar regulatory patterns, several genes experimentally validated to be associated with disease resistance tended to show high TEs in MH63 allele, such as *OsHIR1* (Zhou *et al*., 2010) and *OsLOX3* (Marla and Singh, 2012) (Figure S8A, S8B). MH63 has been reported to be resistant to multiple rice diseases such as blast disease (Xie and Zhang, 2018). Therefore, this similar regulatory pattern is very likely related to the disease resistance phenotype of hybrid SY63 inherited from MH63. Another gene, *OsMH_09G0296900*, was biased to ZS97 at the TE level and identified as a transporter of cytokinin in rice (Figure S8C). The protein encoded by this gene could promote the growth of rice, which might play an important role in the faster growth of ZS97 and SY63 at the early seedling stage compared with MH63 (Xu *et al*., 2004). Pattern B included approximately 40.1% of the total TE-only genes (Figure 5F). Although the number of such TE-only genes was too small for GO enrichment analysis, a large proportion of these genes could still be functionally annotated (Data S10). For example, the gene *OsMH_06G0024600* (*LOC_Os06g03860*) was switched to higher TE in MH63 allele in the root of SY63, which is a phosphate (Pi) transporter involved in maintaining cytosol Pi concentration in rice (Qi and Xiong, 2013) (Figure 5G). Taken together, although the role of allele-specific expression in heterosis has been studied mainly at the transcriptional level, we found that these two patterns only change at the TE level and a large proportion of these genes could be functionally annotated, which might provide a new perspective for studying the role of allele specific expression in heterosis at the translational level.

### Cis-regulatory elements related to divergence of allelic TE

To study the role of *cis*-regulatory elements in determining the divergence of TE-only genes, a total of 361 genes were employed for further analyses. First, we compared the sequence variations between TE-only genes and those without allelic TE divergence (hereafter referred to as control genes; Figure 6). The TE-only genes had more SNPs than the control genes (Figure 6A). Also, a higher SNP density was observed in the TE-only genes (3.15 vs 2.40 SNP/kb; Figure 6B; Wilcoxon test, *P*= 2.43e-05). Besides, in both types of genes, SNPs were enriched in UTR regions (Figure 6C). Compared with the control, the TE-only genes had slightly more SNPs within their 5’UTR regions, but many more SNPs in the CDS and 3’UTR regions. These results implied that variations in these two regions might play a dominant role in determining the allelic TE divergence. In the region near the start codon, we also found more SNPs in TE-only genes (Figure S9B).

**Figure 6.**
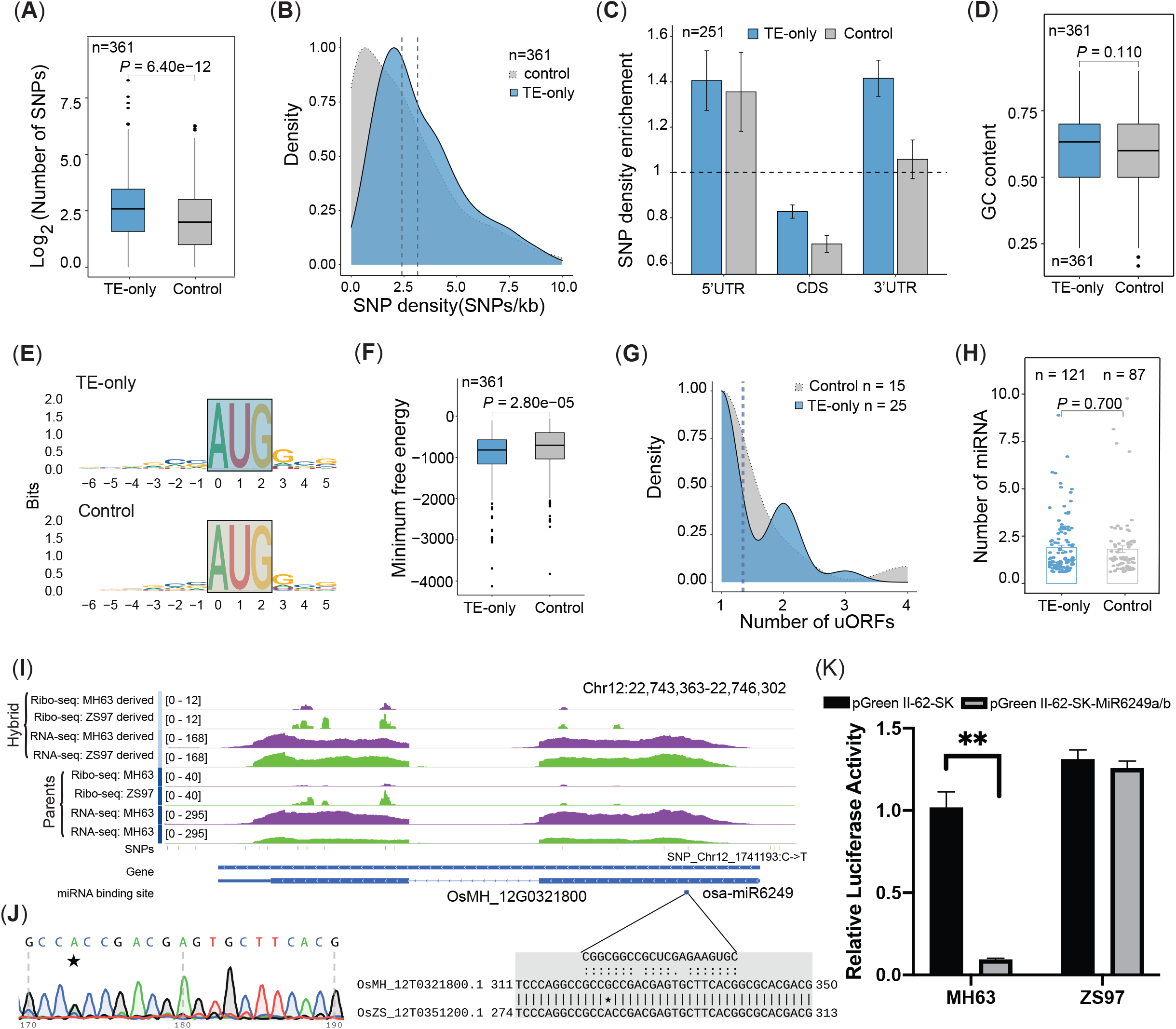
Effect of *cis*-regulatory elements on TE-only genes. TE-only genes and control genes (those without TE divergency between alleles) are compared in (A) Total number of SNPs. (B) SNP density (SNPs/kb). (C) Enrichment of SNPs among three features. Only genes with complete annotation structure (5’UTR, CDS and 3’UTR) were used (n=251). Error bar display ± *s. e. m*. (D) GC content. (E) Kozak sequence around start codon. (F) Minimum free energy (MFE) that reflects the secondary structure of mRNAs. (G) uORFs density. Only genes with uORF were used for comparison (n=25 and 15 for TE-only and control genes, respectively). (H) Number of miRNAs binding site. Only genes with miRNAs binding sites were used for comparison (n=121 and 87 for TE-only and control genes, respectively). Wilcoxon test p-value was labeled in A, D, F, H. (I) Difference in binding site of miRNAs could potentially contribute to the TE divergence in *OsMH_12G0321800*. The Ribo-seq and RNA-seq from phased reads as well as in parents were shown in the first eight tracks followed by SNPs, gene structure and the miRNA bind site tracks. Besides, the mature sequence of osa-miR6249 and target site of MH63 and ZS97 alleles are shown on the bottom panel. Asterisk marked the SNP position within the target site between the two alleles. (J) Sanger sequencing validated the true heterozygous status of the SNP in (I) in hybrid plant SY63. Two peaks with equal height represent adenine (green) and guanine (black). (K) The relative luciferase activity for alleles *OsMH_12G0321800* and *OsZS_12G0351200* in protoplast of hybrid SY63, respectively. The luciferase activity was significantly suppressed in MH63 allele but not in ZS97 allele, indicating that the binding of miRNA osa-miR6249 only happened in the former. Black and grey bar represent control (null vector) and miRNA vector. Error bars display ± *s. d* and p-value ≤ 0.01 was labeled as “**” with student’s t test.

Next, we took several *cis*-regulatory elements into account to study their contribution to the TE divergence of alleles. GC content, as a basic feature, showed no difference between the TE-only genes and control genes (Figure 6D; mean GC content: 0.604 vs. 0.587; Wilcoxon test, *P* = 0.108). The Kozak sequence (Kozak, 1987) was reported to control the translational process in multiple manners (Hata *et al*., 2021). We found that 18 (accounting for 5%) TE-only genes showed variations in their Kozak sequence, while the number was 11 (accounting for 3.05%) for the control genes, showing no significant difference from each other (Fishers exact test, *P* = 0.255). In addition, no significant difference was observed in the position around the start codon as well (Figure 6E). Another important factor that may influence the TE is the secondary structure of mRNAs (Hall *et al*., 1982). We predicted the minimum free energy (MFE) of each mRNA. The results showed that the TE-only genes had a significantly lower MFE than the control genes (Figure 6F; Wilcoxon test, *P*= 2.84e-05), suggesting that the genes with allelic TE divergence tend to possess less stable secondary structure. It has been reported that codon usage could regulate the speed of ribosome movement on mRNAs and thus influence TE (Wang and Roossinck, 2006). We found that the TE-only genes tend to use more optimal codons with high frequencies compared with the control genes (Figure S9C; Wilcoxon test, *P*= 2.98e-04). Moreover, 11.9% of the TE-only genes contained at least one uORF, showing no significant difference from the control genes (7.11%; Fisher exact test, *P* = 0.134). There were also no significant differences in the number of uORFs in these two types of genes (Figure 6G).

Finally, we checked the differences in miRNA binding sites because many previous studies have highlighted the importance of miRNAs in controlling translation in plants (Brodersen *et al*., 2008; Song *et al*., 2019). Between the TE-only and control genes, no significant differences in the number of miRNA binding sites were detected (Figure 6H; Wilcoxon test, *P* = 0.701). However, we still suspected that the binding site divergence within a pair of alleles could contribute to the allelic TE divergence. Therefore, we predicted the genome-wide miRNA binding sites on both MH63 and ZS97 genes. The results showed that 17.5% of the TE-only genes were different between two alleles (Data S11). For example, *OsMH_12G0321800*/*OsZS_12G0351200* contained a binding site for osa-miR6249 only in the CDS region of MH63 allele (Figure 6I). The Ribo-seq signal derived from MH63 allele was significantly lower than that from ZS97 allele. Thus, the binding of this miRNA on MH63 may reduce the translation of this allele but do not affect that of the allele from ZS97. This difference in binding site is likely inherited from their parents, because the phased Ribo-seq signals from the male parent MH63 dataset were also lower than those from the female parent ZS97 dataset (Figure 6I).

Further examination revealed that one SNP (Chr12: 22745704-22745705 in MH63RS3 genome) may be responsible for the difference in binding site between the two alleles (Figure 6I, 6J). To validate this difference, a luciferase reporter assay was performed for the two homologous genes in the protoplast of hybrid SY63 (Methods S4). The transfection of miRNA osa-miR6249 could significantly suppress the luciferase activity of the target gene only in MH63 allele but not in ZS97 allele (Figure 6K, Data S12). Combined with the specific expression of this gene in the root (Figure S9D), it can be speculated that the binding difference of miRNAs may be responsible for the observed TE divergence in these alleles.

## Discussion

Protein synthesis level has long been shown to have poor correlations with mRNA abundance in eukaryotic organisms (Ingolia *et al*., 2009; Battle *et al*., 2015), which may be ascribed to translational regulation to a large extent (Urquidi Camacho *et al*., 2020). In this study, by analysis of the three tissues in two parental varieties and their hybrid, we provide a comprehensive translational regulation profile and allelic TE divergence in rice. First, the high-quality Ribo-seq and RNA-seq data could facilitate investigation of the expression changes at both transcriptional and translational levels. Similar to the findings in animals (Brawand *et al*., 2011; Wang *et al*., 2020), the expression variations across genes at the translational level were generally more significant than those at the transcriptional level due to the extensive translational regulation processes. Different TE distributions were observed in multiple developmental stages, reflecting dynamic translational regulation for various cell types in plants. Besides, although the expression divergence measured by the Euclidean distance varied greatly among different tissues at both the transcriptional and translational levels, the expression distance was generally consistent between the transcriptional and translational levels, suggesting that the evolution of gene regulation is comparable between the two levels. Second, we identified thousands of short translated uORFs and hundreds of potential translated ncORFs from the lncRNAs in rice. These uORFs were somewhat tissue-specific, though most of them were conversed in all examined tissues. Interestingly, we found that not all uORFs repress the translation of mORFs as reported recently in the embryos of *Drosophila melanogaster* (Patraquim *et al*., 2020) and maize(Chotewutmontri and Barkan, 2021), indicating a possibly new mechanism through which uORFs exert their functions. Surprisingly, only a very small proportion of ncORFs were translated as detected in rice. The pipeline used to identify lncRNAs in our previous work has incorporated some tools particularly designed for plants. Thus, it is probably true that only few ncORFs could be translated based on these reliable lncRNAs. Last but most importantly, we conducted a genome-wide survey of ASTE in hybrid rice. Based on multiple tissues and a customized pipeline, we detected hundreds of genes with ASTE, most of which were tissue-specific. Besides, few genes showed coordinated changes at both the transcriptional and translational levels; on the contrary, translational regulation often buffers the changes in gene expression at the transcriptional level, as evidenced by the fact that the number of compensatory genes was approximately ten folds that of reinforcing genes. Apart from the compensatory and reinforcing genes, we also categorized two more types of genes according to allelic expression divergence: mRNA-only genes and TE-only genes. Although there were fewer TE-only genes than mRNA-only genes, the number of TE-only genes was still relatively large. We also observed that certain variations in *cis*-regulatory elements would cause the divergence in TE-only genes, such as the secondary structure of mRNAs, which is consistent with the findings in yeast (Artieri and Fraser, 2014) and mice (Hou *et al*., 2015). Unexpectedly, although the number of uORFs could regulate mORFs either negatively as found in this study or positively as reported in a previous study (Xiong *et al*., 2020) in rice, no significant difference was found in the number of uORFs between TE-only and control genes. The same results were obtained when analyzing the influence of miRNA binding, which has long been shown to inhibit translation by interacting with multiple protein complexes (Sun, 2012; Merchante *et al*., 2017). However, after manual checking, we still found that the divergence in some TE-only genes might be attributable to the binding variations between the two alleles.

Although this study provides many new insights into the translational profile and allelic TE divergence in rice, improvement can still be made in some areas. First, although we identified nearly one hundred uORF-containing TF genes, we failed to find consistent patterns for the functions of their uORFs even after examination of multiple tissues, which has also not been reported in previous studies, particularly in plants. These facts indicate that these uORF-containing TF genes may play diverse roles in different developmental stages of plants. Second, the functions of many lncRNAs have been elucidated in animals especially in humans (Zhang *et al*., 2018); however, there has been no large-scale study in plants. Previous studies in plants have revealed that lncRNAs mainly function through themselves rather than through the production of peptides (Wu *et al*., 2020). This is consistent with our finding that very few peptides from ncORF-containing lncRNAs could be functionally annotated. Degradation of ribosome-bound lncRNAs might be one reason accounting for such phenomenon as revealed in human cells (Carlevaro-Fita *et al*., 2016), which needs to be further validated in plants. Finally, despite that MH63 and ZS97 are two recently selected lines, they exhibited extensive sequence divergence as indicated by the comparative analysis (Zhang *et al*., 2016). In this study, we first selected 30,242 1:1 orthologs accounting for 77.45% of the total non-transposable element genes in MH63RS3 genome, and only 37.54% genes containing SNPs within their CDS were used for the phasing of reads. This means that the reads for more than half of the genes could not be phased to dissect their allelic TE. Besides, understanding the TE divergence of species-specific genes will also be of great significance because the translation of these genes may be involved in many complex biological processes such as disease or stress response due to their various conserved domains (Zhang *et al*., 2016).

In addition to the above-mentioned challenges, future research may be focused on several of the following aspects. First, about a decade ago, Brawand et al. investigated the evolution of gene expression in mammals by using RNA-seq data and found that the evolution rate varies among different lineages and tissues (Brawand *et al*., 2011). Until recently, by using the same materials plus Ribo-seq data, they demonstrated a co-evolution pattern of transcriptome and translatome in mammals (Wang *et al*., 2020). One important reason for their success of study is the innovation of algorithms for measuring the evolution of gene expression at both transcriptional and translational levels. So far, Ribo-seq and corresponding RNA-seq data have become available in several important monocots and dicots such as *Arabidopsis* (Juntawong *et al*., 2014; Hsu *et al*., 2016; Bazin *et al*., 2017), maize (Lei *et al*., 2015), wheat (Guo *et al*., 2015), soybean (Shamimuzzaman and Vodkin, 2018), potato (Wu *et al*., 2019), and rice (Xiong *et al*., 2020), providing valuable resources for studying the evolution of genes at the transcriptional and translational levels in plants (Voelckel *et al*., 2017). However, in the long history after the divergence between animals and plants, plants have acquired many distinct features totally different from those of animals in both genome and morphology. Hence, more new and robust algorithms are required to dissect the evolution history of plants at the translational level. Second, when evaluating the impact of *cis*-regulatory elements on the allelic TE divergence, we mainly considered some mRNA sequence features such as uORFs within 5’UTR and miRNA binding sites. However, we ignored another important factor, the epigenetic modification on mRNAs such as N6-methyladenosine (m6A) methylation, which is currently a fascinating research hotspot in the mRNA studies of plants. Many studies of mammals have revealed that m6A modification can mediate multiple biological processes such as mRNA degradation, stability and most importantly translation (Visvanathan and Somasundaram, 2018). However, such studies are still relatively rare in plants, and most of the available studies are mainly focused on the roles of m6A modification in plant development (Růžička *et al*., 2017) and stress response (Scutenaire *et al*., 2018). Thus, whether these modifications of mRNAs also contribute to allelic TE divergence remains to be explored. Besides, some small RNAs like small interfering RNAs (siRNAs) may also contribute to the TE divergence in alleles in a way similar to that of miRNAs (Brodersen *et al*., 2008). Last but not least, as one of the most complex issues in plant biological research, heterosis has been attracting attention for more than one hundred years due to its great significance in crop breeding (Schnable and Springer, 2013). With the availability of methods to quantify the gene expression level, such as gene expression microarrays and RNA-seq, breeders can validate many modes of gene action such as additivity and under- or over-dominance (Hochholdinger and Hoecker, 2007). However, very little research has been conducted on allele-specific expression at the translational level (Zhu *et al*., 2021). The ignorance of heterosis at the translational level in previous studies is probably due to the intrinsic complexity of heterosis, as the observed heterosis can be hardly explained by one single mechanism. Despite such great difficulties, we still observed two major regulatory patterns at the TE level, which account for more than 80% of the TE-only genes, by comparing the variations at both transcriptional and translational levels between the parents and hybrid (Figure 5F). These two patterns only showed divergence at the TE level, highlighting the potential important role of translational regulation in heterosis, which can be hardly inferred by only using RNA-seq data. More innovative experimental methods may be developed to help explore this phenomenon in more detail, which can enhance our understanding of the mechanism for heterosis and finally facilitate crop breeding for food safety.

In summary, by combining Ribo-seq and RNA-seq data from three tissues of three rice varieties, we obtained a comprehensive translational profile that highlights the variations of translational regulation among tissues. Besides, by taking advantages of the hybrid and its parents, this study dissected the allelic TE divergence in hybrid plants, which may be of great significance for expanding the knowledge of crop breeding.

## Materials and Methods

### Plant materials, library construction and routine data process

The seeds of three rice varieties, MH63, ZS97 and SY63, were first germinated by soaking in sterile water for two days, and then transferred to the light incubator for growth under normal conditions (28°C for 14 h in light and 26°C for 10 h in dark, 70% humidity) in culture solution (Yoshida, 1976). The samples of young leaves and roots at the four-leaf stage and panicles were collected for both RNA-seq and Ribo-seq library construction. All fresh samples were immediately frozen in −80°C liquid nitrogen until use. The RNA-seq library construction and sequencing were performed as described in our early work (Zhou *et al*., 2021). The Ribo-seq library construction was carried out according to a previous study (Ingolia *et al*., 2012), and more details are described in Methods S1. Multiple routine tools were used to process the RNA-seq and Ribo-seq data and the details can be found in Methods S2.

### Analysis of unique features of Ribo-seq data

The general features such as length and genome distribution of RPFs were calculated using custom R scripts. The P-site offset for each read length was determined with psite command in Plastid (v0.4.8) (Dunn and Weissman, 2016), and RPFs with length 25–31nt were used for subsequent analysis. The 15^th^ base from the 5’ end of reads was considered as a simulative P-site offset while using RNA-seq as the control. Then, riboWaltz (Lauria *et al*., 2018) was used to calculate the distribution of P-site signal among 5’UTR, CDS and 3’UTR. For the analysis of 3-nucleotide periodicity, we first summed up the P-site across every position of all mRNAs and then normalized the data with the average value of the upstream 20 bases from the start codon. The relative codon usage in Figure S1D, S6B, S9C was calculated in CodonW (v1.4.2, http://codonw.sourceforge.net/) with respective genes.

### Calculation of TE and expression distance

The TE for each gene with the TPM value >= 0.1 at both levels was defined as the ratio of TPM_Ribo-seq_ to TPM_RNA-seq_, and the TE range across genes in each sample was calculated as the ratio of 97.5^th^ to 2.5^th^ percentile of TEs to eliminate the effects of outliers. The expression variances were calculated from the expression matrix in R (Team, 2013), and the divergence in expression profile (expression distance) between a pair of tissues was measured by the Euclidean distance (Glazko and Mushegian, 2010):

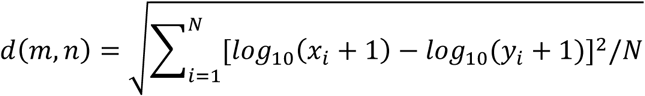

In the equation, *m, n* refer to sample *m* and *n*; the variables *x* and *y* are the TPM values for gene *i* at either transcriptional or translational level in sample *m* and *n*, respectively; the variable *N* represents the number of transcribed genes (*N* = 23,957) in all three varieties. The larger *d(m, n)* indicates a greater variation in expression profile.

### Identification of uORFs

The uORFs longer than 10 aa were identified with an AUG start codon followed by an in-frame stop codon within the annotated 5’ UTR of primary protein-coding mRNA in MH63RS3. ORFs completely contained within another were discarded. Then, the sequenced reads mapped to uORFs were counted with featureCounts (v2.0.0) (Liao *et al*., 2014). ORFs with RPFs greater than three in any replicates were considered as active uORFs in each sample. In addition, we downloaded all rice datasets of TFs from PlantTFDB (v5.0) (Tian *et al*., 2020) to analyze the TF types of uORF-containing genes.

### Detection and de novo functional annotation of actively translated ORFs in lncRNAs

The detection of actively translated ORFs was performed using RiboTaper (v1.3.1a)(Calviello *et al*., 2016) with the pairing RPFs length and P-site offset: 25, 26, 27, 28, 29, 30 and 31 and 9, 10, 11, 12, 13, 13 and 13. The ORFs with the same stop codon but different start codon was regarded as the same ORF across samples. The lncRNAs in MH63RS3 genome were re-identified with the same pipeline described in our previous work (Zhou *et al*., 2021). Three strategies were used for the *de novo* functional annotation of active ORFs. First, each putative active ORF was queried against multiple databases through a local InterProScan 5 (v5.48-83.0) (Jones *et al*., 2014) with a hit e-value threshold of 1e-5. Second, all annotated protein sequences in MH63RS3 with lengths shorter than 100 aa were treated as the set of known small proteins. The active ORFs were queried against this database through BLASTP (v2.9.0+) (Camacho *et al*., 2009) with an e-value threshold of 1e-5. Third, TMHMM 2 (Krogh *et al*., 2001) and SignalP-5.0 (Almagro Armenteros *et al*., 2019) with default parameters were used to predict transmembrane proteins and signal peptides. sORFs were first selected from protein-coding genes with CDS shorter than 300nt and only those with translational activity predicted by RiboTaper were retained. The control sets were randomly selected from actively translated genes with CDS length longer than 300nt, and the number was the same as that of the above identified sORFs.

### Read phasing and identification of genes with ASTE

To identify genes with ASTE, it is necessary to distinguish reads derived from different parents. Previous studies in yeast (Artieri and Fraser, 2014) or mice (Hou *et al*., 2015) adopted the perfect-matching strategy for read phasing, in which the reads should be perfectly matched to the SNP position of either homologous genes; however, the perfectly matched reliable reads in one parent should have a mismatch in the SNP position of another parent. Thus, we used PP2PG (v1.0) (Feng *et al*., 2021) which implements such algorithm to phase the reads of SY63. Before the reads were phased (Figure 5A), nucmer, delta-filter and show-snps in MUMmer4 packages (v4.0.0beta2) (Marçais *et al*., 2018) were used to call SNPs between MH63RS3 and ZS97RS3 genomes and a total of 1,757,026 SNPs were identified. In order to evaluate the phasing quality, we also applied the same procedure to the Ribo-seq and RNA-seq reads from both MH63 and ZS97. The error rate, which benchmarks the proportion of reads in MH63 datasets that were wrongly assigned to ZS97 or vice versa, was defined as ZS97-derived reads / (MH63-derived reads + ZS97-derived reads) in MH63 datasets or MH63-derived reads / (MH63-derived reads + ZS97-derived reads) in ZS97 datasets. Another quality index, separation rate, was used to describe how many reads in the total datasets could be phased and was calculated as (MH63-derived reads + ZS97-derived reads) / (MH63-derived reads + ZS97-derived reads + Unknown reads). The phasing results are listed in Table S4 and Data S5.

After acquiring the phased reads, by using the intersect subcommand in bedtools (v2.28.0) (Quinlan and Hall, 2010), we first selected 14,659 genes from all 1:1 orthologs between MH63 and ZS97 genomes that contained SNPs within their CDS region and thus could be used to determine the allelic origin of reads mapped on them. Next featureCounts (v2.0.0) (Liao *et al*., 2014) was used to assign the phased reads to these genes and two read count matrices could thus be constructed at the transcriptional and translational levels. Finally, we modified the R scripts in deltaTE (Chothani *et al*., 2019) and used DESeq2 (Love *et al*., 2014) to identify the genes with significant differences in transcription as well as the TE levels under the FDR<=0.05 and |log_2_(fold change)|>=1 cut-off. According to the values of log_2_ (fold change) at the transcriptional and TE levels, these genes with allelic expression divergence were further divided into four categories: the genes whose expression changed only at the mRNA level (mRNA-only genes), only at the TE level (TE-only genes), compensatory genes and reinforcing genes. All the above analyses were performed in a custom R script.

### Identification of cis-regulatory elements

Several mRNA sequence features were identified in this study. First, the SNPs called in the previous step were transferred from the genomic position to the mRNA position by R package GenomicFeatures (v1.40.1) (Lawrence *et al*., 2013). To reveal the unique features of TE-only genes, we randomly selected an equal number of homologous genes without TE divergence in a pair of alleles as the control genes. The SNP number, enriched density, distribution in different features and the density before the start codon were calculated using the custom R script. Second, the GC content of 30 bases around the start codon (–15, +14 bases) and Kozak sequence (defined as the sequence from –6 to +5 bases relative to the start codon) for each involved gene was calculated or extracted using bedtools. Third, RNAfold (v 2.4.16) (Lorenz *et al*., 2011) was used to calculate the minimum free energy (MFE) around the start codon (–15, +14 bases) with default parameters at a temperature of 28°C. Finally, to obtain highly reliable miRNA binding sites, we downloaded all the mature rice miRNA sequences from miRBase (release 22)(Kozomara *et al*., 2019) and then only the binding sites predicted by both TargetFinder (https://github.com/carringtonlab/TargetFinder) and psRNAtarget (2017) (Dai *et al*., 2018) were retained for further comparison analysis. The TE-only and control genes from parental MH63 were used for the comparison in Fig 6D-H.

### GO enrichment analysis

We improved the functional annotation of genes for MH63RS3 genome (Methods S3), and the script run_GOseq.pl in Trinity packages (v2.8.5)(Haas *et al*., 2013) was used for GO enrichment analysis.

### Statistical methods

In this study, Fisher exact test, Wilcoxon test, Kolmogorov-Smirnov test and Pearson correlation coefficient calculation were performed by fisher.test, wilcox.test, ks.test and cor.test in R, respectively. Additional data processing and plotting methods are described in Methods S5.

## Data availability

The RNA-seq data have been deposited at National Genomics Data Center (CRA002886) and the Ribo-seq data have been submitted to the GenBank database under the accession number PRJNA725700. The custom R scripts used in this study are freely available from GitHub at https://github.com/Zhuxitong/Rice_Ribo-seq.

## Acknowledgements

This work was supported by the National Natural Science Foundation of China (31871269), Hubei Provincial Natural Science Foundation of China (2019CFA014) and the starting research grant for High-level Talents from Guangxi University. The computations in this study were run on the bioinformatics computing platform of the National Key Laboratory of Crop Genetic Improvement, Huazhong Agricultural University.

## Author contributions

L.-L.C., J.-X.G., X.-T.Z., R. Z. designed the project, X.-T.Z., R. Z., Y.-Y.Z. and J.-W.F. analyzed the data; X.-T.Z. R. Z. and J. C. performed experiments; X.-T.Z., R. Z. M.T.Q., J.-W.Z., J.-X.G. and L.-L.C. wrote and revised the paper.

## Supporting Information

**Figure S1** Data quality of Ribo-seq.

**Figure S2** Global expression changes among different varieties and tissues.

**Figure S3** Correlation of expression value between RNA-seq and Ribo-seq.

**Figure S4** uORF-mediated effects on translation efficiency.

**Figure S5** Characteristics of translated lncRNAs.

**Figure S6** The comparison of TE and codon usage among different kind of ORFs.

**Figure S7** Characteristics of genes with ASTE.

**Figure S8** RNA-seq and Ribo-seq signals for three cloned genes with expression bias.

**Figure S9** Comparison of TE-only and control genes and expression of gene *OsMH_12G0321800*.

**Table S1** Transcribed and translated genes in each sample (TPM >=0.1).

**Table S2** Number of uORFs (uORFs-contained genes) in each sample.

**Table S3** Number of sORFs in lncRNAs / protein-coding genes in each sample.

**Table S4** Phasing summary.

**Data S1**: Summary of data quality for Ribo-seq and RNA-seq data.

**Data S2**: GO enrichment results for genes with top and bottom 5% TEs in each sample.

**Data S3**: List of uORFs and GO enrichment results for uORFs-contained genes.

**Data S4**: List of ncORFs and their predicted functions.

**Data S5**: Phasing summary for Ribo-seq and RNA-seq in SY63.

**Data S6:** DESeq2 results for genes with ASTE in three tissues of SY63.

**Data S7:** RNA-seq and Ribo-seq phased reads count matrix in SY63 and parents.

**Data S8**: List of four categories genes with allelic expression divergence in each tissue of SY63.

**Data S9**: GO enrichment results for mRNA-only and TE-only genes.

**Data S10:** Regulatory pattern and function annotation for TE-only genes in three tissues of SY63.

**Data S11**: List of alleles with difference in binding site of miRNAs.

**Data S12**: Signals from luciferase reporter assay for osa-miR6249 in hybrid protoplast.

**Methods S1** Ribo-seq library construction and sequencing.

**Methods S2** Routine data process of RNA-seq and Ribo-seq data.

**Methods S3** Improvement of functional annotation for protein-coding genes in MH63RS3 genome.

**Methods S4** Luciferase reporter assay.

**Methods S5** Statistical test, data process and plotting methods.

## Parsed Citations

Almagro Armenteros JJ, Tsirigos KD, Sønderby CK, Petersen TN, Winther O, Brunak S, von Heijne G, Nielsen H (2019) SignalP 5.0 improves signal peptide predictions using deep neural networks. Nat Biotechnol 37:420–423

Artieri CG, Fraser HB (2014) Evolution at two levels of gene expression in yeast. Genome research 24:411–421

Battle A, Khan Z, Wang SH, Mitrano A, Ford MJ, Pritchard JK, Gilad Y (2015) Impact of regulatory variation from RNA to protein. Science 347:664

Bazin J, Baerenfaller K, Gosai SJ, Gregory BD, Crespi M, Bailey-Serres J (2017) Global analysis of ribosome-associated noncoding RNAs unveils new modes of translational regulation. Proceedings of the National Academy of Sciences 114:E10018

Brar GA, Weissman JS (2015) Ribosome profiling reveals the what, when, where and how of protein synthesis. Nature Reviews Molecular Cell Biology 16:651–664

Brawand D, Soumillon M, Necsulea A, Julien P, Csárdi G, Harrigan P, Weier M, Liechti A, Aximu-Petri A, Kircher M, Albert FW, Zeller U, Khaitovich P, Grützner F, Bergmann S, Nielsen R, Pääbo S, Kaessmann H (2011) The evolution of gene expression levels in mammalian organs. Nature 478:343–348

Brodersen P, Sakvarelidze-Achard L, Bruun-Rasmussen M, Dunoyer P, Yamamoto YY, Sieburth L, Voinnet O (2008) Widespread Translational Inhibition by Plant miRNAs and siRNAs. Science 320:1185

Calviello L, Mukherjee N, Wyler E, Zauber H, Hirsekorn A, Selbach M, Landthaler M, Obermayer B, Ohler U (2016) Detecting actively translated open reading frames in ribosome profiling data. Nat Methods 13:165–170

Calvo SE, Pagliarini DJ, Mootha VK (2009) Upstream open reading frames cause widespread reduction of protein expression and are polymorphic among humans. Proceedings of the National Academy of Sciences 106:7507–7512

Camacho C, Coulouris G, Avagyan V, Ma N, Papadopoulos J, Bealer K, Madden TL (2009) BLAST+: architecture and applications. BMC Bioinformatics 10:421

Cardoso-Moreira M, Halbert J, Valloton D, Velten B, Chen C, Shao Y, Liechti A, Ascenção K, Rummel C, Ovchinnikova S (2019) Gene expression across mammalian organ development. Nature 571:505–509

Carlevaro-Fita J, Rahim A, Guigó R, Vardy LA, Johnson R (2016) Cytoplasmic long noncoding RNAs are frequently bound to and degraded at ribosomes in human cells. Rna 22:867–882

Chotewutmontri P, Barkan A (2021) Ribosome profiling elucidates differential gene expression in bundle sheath and mesophyll cells in maize. Plant Physiology 187:59–72

Chothani S, Adami E, Ouyang JF, Viswanathan S, Hubner N, Cook SA, Schafer S, Rackham OJL (2019) deltaTE: Detection of Translationally Regulated Genes by Integrative Analysis of Ribo-seq and RNA-seq Data. Current Protocols in Molecular Biology 129:e108

Clark JW, Donoghue PC (2018) Whole-genome duplication and plant macroevolution. Trends in plant science 23:933–945

Crick F (1970) Central Dogma of Molecular Biology. Nature 227:561–563

Csárdi G, Franks A, Choi DS, Airoldi EM, Drummond DA (2015) Accounting for experimental noise reveals that mRNA levels, amplified by post-transcriptional processes, largely determine steady-state protein levels in yeast. PLoS Genet 11:e1005206

Dai X, Zhuang Z, Zhao PX (2018) psRNATarget: a plant small RNA target analysis server (2017 release). Nucleic acids research 46:W49–W54

Dunn JG, Foo CK, Belletier NG, Gavis ER, Weissman JS (2013) Ribosome profiling reveals pervasive and regulated stop codon readthrough in Drosophila melanogaster. Elife 2:e01179

Dunn JG, Weissman JS (2016) Plastid: nucleotide-resolution analysis of next-generation sequencing and genomics data. BMC Genomics 17:958

Feng J-W, Lu Y, Shao L, Zhang J, Li H, Chen L-L (2021) Phasing analysis of the transcriptome and epigenome in a rice hybrid reveals the inheritance and difference in DNA methylation and allelic transcription regulation. Plant Communications: 100185

Fields Alexander P, Rodriguez Edwin H, Jovanovic M, Stern-Ginossar N, Haas Brian J, Mertins P, Raychowdhury R, Hacohen N, Carr Steven A, Ingolia Nicholas T, Regev A, Weissman Jonathan S (2015) A Regression-Based Analysis of Ribosome-Profiling Data Reveals a Conserved Complexity to Mammalian Translation. Molecular Cell 60:816–827

Glazko G, Mushegian A (2010) Measuring gene expression divergence: the distance to keep. Biology Direct 5:51

Guo W, Zhang J, Zhang N, Xin M, Peng H, Hu Z, Ni Z, Du J (2015) The wheat NAC transcription factor TaNAC2L is regulated at the transcriptional and post-translational levels and promotes heat stress tolerance in transgenic Arabidopsis. PLoS One 10:e0135667

Haas BJ, Papanicolaou A, Yassour M, Grabherr M, Blood PD, Bowden J, Couger MB, Eccles D, Li B, Lieber M, MacManes MD, Ott M, Orvis J, Pochet N, Strozzi F, Weeks N, Westerman R, William T, Dewey CN, Henschel R, LeDuc RD, Friedman N, Regev A (2013) De novo transcript sequence reconstruction from RNA-seq using the Trinity platform for reference generation and analysis. Nature Protocols 8:1494–1512

Hall MN, Gabay J, Débarbouillé M, Schwartz M (1982) A role for mRNA secondary structure in the control of translation initiation. Nature 295:616–618

Hata T, Satoh S, Takada N, Matsuo M, Obokata J (2021) Kozak Sequence Acts as a Negative Regulator for De Novo Transcription Initiation of Newborn Coding Sequences in the Plant Genome. Molecular Biology and Evolution 38:2791–2803

Hochholdinger F, Hoecker N (2007) Towards the molecular basis of heterosis. Trends in Plant Science 12:427–432

Hou J, Wang X, McShane E, Zauber H, Sun W, Selbach M, Chen W (2015) Extensive allele-specific translational regulation in hybrid mice. Molecular Systems Biology 11:825

Hsu PY, Calviello L, Wu H-YL, Li F-W, Rothfels CJ, Ohler U, Benfey PN (2016) Super-resolution ribosome profiling reveals unannotated translation events in Arabidopsis. Proceedings of the National Academy of Sciences 113:E7126

Ingolia NT (2014) Ribosome profiling: new views of translation, from single codons to genome scale. Nature Reviews Genetics 15:205–213

Ingolia NT, Brar GA, Rouskin S, McGeachy AM, Weissman JS (2012) The ribosome profiling strategy for monitoring translation in vivo by deep sequencing of ribosome-protected mRNA fragments. Nature Protocols 7:1534–1550

Ingolia NT, Ghaemmaghami S, Newman JRS, Weissman JS (2009) Genome-Wide Analysis in Vivo of Translation with Nucleotide Resolution Using Ribosome Profiling. Science 324:218

Jones P, Binns D, Chang H-Y, Fraser M, Li W, McAnulla C, McWilliam H, Maslen J, Mitchell A, Nuka G, Pesseat S, Quinn AF, Sangrador-Vegas A, Scheremetjew M, Yong S-Y, Lopez R, Hunter S (2014) InterProScan 5:genome-scale protein function classification. Bioinformatics (Oxford, England) 30:1236–1240

Juntawong P, Girke T, Bazin J, Bailey-Serres J (2014) Translational dynamics revealed by genome-wide profiling of ribosome footprints in Arabidopsis. Proceedings of the National Academy of Sciences 111:E203–E212

Kang K, Shim Y, Gi E, An G, Paek N-C (2019) Mutation of ONAC096 Enhances Grain Yield by Increasing Panicle Number and Delaying Leaf Senescence during Grain Filling in Rice. International journal of molecular sciences 20:5241

Klepikova AV, Penin AA (2019) Gene Expression Maps in Plants: Current State and Prospects. Plants 8:309

Kozak M (1987) An analysis of 5’-noncoding sequences from 699 vertebrate messenger RNAs. Nucleic acids research 15:8125– 8148

Kozomara A, Birgaoanu M, Griffiths-Jones S (2019) miRBase: from microRNA sequences to function. Nucleic Acids Research 47:D155–D162

Krogh A, Larsson B, Von Heijne G, Sonnhammer EL (2001) Predicting transmembrane protein topology with a hidden Markov model: application to complete genomes. Journal of molecular biology 305:567–580

Lauria F, Tebaldi T, Bernabò P, Groen EJ, Gillingwater TH, Viero G (2018) riboWaltz: Optimization of ribosome P-site positioning in ribosome profiling data. PLoS computational biology 14:e1006169

Lawrence M, Huber W, Pages H, Aboyoun P, Carlson M, Gentleman R, Morgan MT, Carey VJ (2013) Software for computing and annotating genomic ranges. PLoS Comput Biol 9:e1003118

Lei L, Shi J, Chen J, Zhang M, Sun S, Xie S, Li X, Zeng B, Peng L, Hauck A, Zhao H, Song W, Fan Z, Lai J (2015) Ribosome profiling reveals dynamic translational landscape in maize seedlings under drought stress. The Plant Journal 84:1206–1218

Liao Y, Smyth GK, Shi W (2014) featureCounts: an efficient general purpose program for assigning sequence reads to genomic features. Bioinformatics 30:923–930

Liu M-J, Wu S-H, Wu J-F, Lin W-D, Wu Y-C, Tsai T-Y, Tsai H-L, Wu S-H (2013) Translational Landscape of Photomorphogenic Arabidopsis. The Plant Cell 25:3699

Lorenz R, Bernhart SH, Zu Siederdissen CH, Tafer H, Flamm C, Stadler PF, Hofacker IL (2011) ViennaRNA Package 2.0. Algorithms for molecular biology 6:1–14

Love MI, Huber W, Anders S (2014) Moderated estimation of fold change and dispersion for RNA-seq data with DESeq2. Genome Biology 15:550

Marçais G, Delcher AL, Phillippy AM, Coston R, Salzberg SL, Zimin A (2018) MUMmer4: A fast and versatile genome alignment system. PLOS Computational Biology 14:e1005944

Marla SS, Singh V (2012) LOX genes in blast fungus (Magnaporthe grisea) resistance in rice. Functional & integrative genomics 12:265–275

Matsubayashi Y (2014) Posttranslationally Modified Small-Peptide Signals in Plants. Annual Review of Plant Biology 65:385–413

McManus CJ, May GE, Spealman P, Shteyman A (2014) Ribosome profiling reveals post-transcriptional buffering of divergent gene expression in yeast. Genome research 24:422–430

Merchante C, Stepanova AN, Alonso JM (2017) Translation regulation in plants: an interesting past, an exciting present and a promising future. The Plant Journal 90:628–653

Muzzey D, Sherlock G, Weissman JS (2014) Extensive and coordinated control of allele-specific expression by both transcription and translation in Candida albicans. Genome Res 24:963–973

Noor Z, Ahn SB, Baker MS, Ranganathan S, Mohamedali A (2021) Mass spectrometry–based protein identification in proteomics-a review. Briefings in Bioinformatics 22:1620–1638

Pajerowska-Mukhtar KM, Wang W, Tada Y, Oka N, Tucker CL, Fonseca JP, Dong X (2012) The HSF-like transcription factor TBF1 is a major molecular switch for plant growth-to-defense transition. Current Biology 22:103–112

Patraquim P, Mumtaz MAS, Pueyo JI, Aspden JL, Couso J-P (2020) Developmental regulation of canonical and small ORF translation from mRNAs. Genome Biology 21:128

Puranik S, Sahu PP, Srivastava PS, Prasad M (2012) NAC proteins: regulation and role in stress tolerance. Trends in Plant Science 17:369–381

Qi Z, Xiong L (2013) Characterization of a Purine Permease Family Gene Os PUP 7 Involved in Growth and Development Control in Rice. Journal of integrative plant biology 55:1119–1135

Quinlan AR, Hall IM (2010) BEDTools: a flexible suite of utilities for comparing genomic features. Bioinformatics 26:841–842

Ribone PA, Capella M, Arce AL, Chan RL (2017) A uORF Represses the Transcription Factor AtHB1 in Aerial Tissues to Avoid a Deleterious Phenotype. Plant Physiology 175:1238

Růžička K, Zhang M, Campilho A, Bodi Z, Kashif M, Saleh M, Eeckhout D, El-Showk S, Li H, Zhong S (2017) Identification of factors required for m6A mRNA methylation in Arabidopsis reveals a role for the conserved E3 ubiquitin ligase HAKAI. New Phytologist 215:157–172

Schnable PS, Springer NM (2013) Progress toward understanding heterosis in crop plants. Annual review of plant biology 64:71– 88

Scutenaire J, Deragon J-M, Jean V, Benhamed M, Raynaud C, Favory J-J, Merret R, Bousquet-Antonelli C (2018) The YTH domain protein ECT2 is an m6A reader required for normal trichome branching in Arabidopsis. The Plant Cell 30:986–1005

Shamimuzzaman M, Vodkin L (2018) Ribosome profiling reveals changes in translational status of soybean transcripts during immature cotyledon development. PLOS ONE 13:e0194596

Song J-M, Xie W-Z, Wang S, Guo Y-X, Koo D-H, Kudrna D, Gong C, Huang Y, Feng J-W, Zhang W, Zhou Y, Zuccolo A, Long E, Lee S, Talag J, Zhou R, Zhu X-T, Yuan D, Udall J, Xie W, Wing RA, Zhang Q, Poland J, Zhang J, Chen L-L (2021) Two gap-free reference genomes and a global view of the centromere architecture in rice. Molecular Plant 14:1757–1767

Song X, Li Y, Cao X, Qi Y (2019) MicroRNAs and Their Regulatory Roles in Plant–Environment Interactions. Annual Review of Plant Biology 70:489–525

Sun G (2012) MicroRNAs and their diverse functions in plants. Plant molecular biology 80:17–36

Team RC (2013) R: A language and environment for statistical computing.

Tian F, Yang D-C, Meng Y-Q, Jin J, Gao G (2020) PlantRegMap: charting functional regulatory maps in plants. Nucleic Acids Research 48:D1104–D1113

Urquidi Camacho RA, Lokdarshi A, von Arnim AG (2020) Translational gene regulation in plants: A green new deal. WIREs RNA 11:e1597

Visvanathan A, Somasundaram K (2018) mRNA Traffic Control Reviewed: N6-Methyladenosine (m6A) Takes the Driver’s Seat. Bioessays 40:1700093

Voelckel C, Gruenheit N, Lockhart P (2017) Evolutionary Transcriptomics and Proteomics: Insight into Plant Adaptation. Trends in Plant Science 22:462–471

Wang L, Roossinck MJ (2006) Comparative analysis of expressed sequences reveals a conserved pattern of optimal codon usage in plants. Plant Mol Biol 61:699–710

Wang Z-Y, Leushkin E, Liechti A, Ovchinnikova S, Mößinger K, Brüning T, Rummel C, Grützner F, Cardoso-Moreira M, Janich P, Gatfield D, Diagouraga B, de Massy B, Gill ME, Peters AHFM, Anders S, Kaessmann H (2020) Transcriptome and translatome co-evolution in mammals. Nature 588:642–647

Weinberg DE, Shah P, Eichhorn SW, Hussmann JA, Plotkin JB, Bartel DP (2016) Improved Ribosome-Footprint and mRNA Measurements Provide Insights into Dynamics and Regulation of Yeast Translation. Cell Reports 14:1787–1799

Wu H-YL, Song G, Walley JW, Hsu PY (2019) The Tomato Translational Landscape Revealed by Transcriptome Assembly and Ribosome Profiling. Plant Physiology 181:367

Wu L, Liu S, Qi H, Cai H, Xu M (2020) Research Progress on Plant Long Non-Coding RNA. Plants 9:408

Xie F, Zhang J (2018) Shanyou 63:an elite mega rice hybrid in China. Rice (New York, N.Y.) 11:17–17

Xiong Q, Zhong L, Du J, Zhu C, Peng X, He X, Fu J, Ouyang L, Bian J, Hu L, Sun X, Xu J, Zhou D, Cai Y, Fu H, He H, Chen X (2020) Ribosome profiling reveals the effects of nitrogen application translational regulation of yield recovery after abrupt drought-flood alternation in rice. Plant Physiology and Biochemistry 155:42–58

Xu CG, Li XQ, Xue Y, Huang YW, Gao J, Xing YZ (2004) Comparison of quantitative trait loci controlling seedling characteristics at two seedling stages using rice recombinant inbred lines. Theor Appl Genet 109:640–647

Yang X, Cui J, Song B, Yu Y, Mo B, Liu L (2020) Construction of High-Quality Rice Ribosome Footprint Library. Frontiers in Plant Science 11:1381

Yoshida S (1976) Routine procedure for growing rice plants in culture solution. Laboratory manual for physiological studies of rice: 61–66

Zhang J, Chen L-L, Xing F, Kudrna DA, Yao W, Copetti D, Mu T, Li W, Song J-M, Xie W, Lee S, Talag J, Shao L, An Y, Zhang C-L, Ouyang Y, Sun S, Jiao W-B, Lv F, Du B, Luo M, Maldonado CE, Goicoechea JL, Xiong L, Wu C, Xing Y, Zhou D-X, Yu S, Zhao Y, Wang G, Yu Y, Luo Y, Zhou Z-W, Hurtado BEP, Danowitz A, Wing RA, Zhang Q (2016) Extensive sequence divergence between the reference genomes of two elite indica rice varieties Zhenshan 97 and Minghui 63. Proceedings of the National Academy of Sciences of the United States of America 113:E5163–E5171

Zhang Y, Tao Y, Liao Q (2018) Long noncoding RNA: a crosslink in biological regulatory network. Briefings in Bioinformatics 19:930–945

Zhao D, Hamilton JP, Hardigan M, Yin D, He T, Vaillancourt B, Reynoso M, Pauluzzi G, Funkhouser S, Cui Y, Bailey-Serres J, Jiang J, Buell CR, Jiang N (2017) Analysis of Ribosome-Associated mRNAs in Rice Reveals the Importance of Transcript Size and GC Content in Translation. G3 (Bethesda, Md.) 7:203–219

Zhou L, Cheung M-Y, Li M-W, Fu Y, Sun Z, Sun S-M, Lam H-M (2010) Rice hypersensitive induced reaction protein 1 (OsHIR1) associates with plasma membrane and triggers hypersensitive cell death. BMC plant biology 10:1–10

Zhou R, Sanz-Jimenez P, Zhu X-T, Feng J-W, Shao L, Song J-M, Chen L-L (2021) Analysis of Rice Transcriptome Reveals the LncRNA/CircRNA Regulation in Tissue Development. Rice 14:14

Zhu W, Chen S, Zhang T, Qian J, Luo Z, Zhao H, Zhang Y, Li L (2021) Dynamic patterns of the translatome in a hybrid triplet show translational fractionation of the maize subgenomes. The Crop Journal

